# GETgene-AI: A Framework for Prioritizing Actionable Cancer Drug Targets

**DOI:** 10.1101/2025.01.21.634201

**Authors:** Adrian Gu, Jake Y Chen

## Abstract

Prioritizing actionable drug targets is a critical challenge in cancer research, where high-dimensional genomic data and the complexity of tumor biology often hinder effective prioritization. To address this, we developed **GETgene-AI**, a novel computational framework that integrates network-based prioritization, machine learning, and automated literature analysis to prioritize and rank potential therapeutic targets. Central to GETgene-AI is the **G.E.T. strategy**, which combines three data streams: mutational frequency (G List), differential expression (E List), and known drug targets (T List). These components are iteratively refined and ranked using the **Biological Entity Expansion and Ranking Engine (BEERE)**, leveraging protein-protein interaction networks, functional annotations, and experimental evidence. Additionally, GETgene-AI incorporates GPT-4o, an advanced large language model, to automate literature-based ranking, reducing manual curation and increasing efficiency.

In this study, we applied GETgene-AI to pancreatic cancer as a case study. The framework successfully prioritized high-priority targets such as **PIK3CA** and **PRKCA**, validated through experimental evidence and clinical relevance. Benchmarking against GEO2R and STRING demonstrated GETgene-AI’s superior performance, achieving higher precision, recall, and efficiency in prioritizing actionable targets. Moreover, the framework mitigated false positives by deprioritizing genes lacking functional or clinical significance.

While demonstrated on pancreatic cancer, the modular design of GETgene-AI enables scalability across diverse cancers and diseases. By integrating multi-omics datasets with advanced computational and AI-driven approaches, GETgene-AI provides a versatile and robust platform for accelerating cancer drug discovery. This framework bridges computational innovations with translational research to improve patient outcomes.

## INTRODUCTION

Traditional chemotherapeutic agents, which non-specifically target rapidly dividing cells (Gu et al., 2023; Sun et al., 2021), are contested with the promise of targeted therapies that disrupt specific molecular pathways governing cell survival and apoptosis (Lim et al., 2019; Sellers & Fisher, 1999). Drug target discovery is pivotal for advancing cancer therapies, yet traditional approaches face three critical limitations. First, manual curation of literature and static biomedical databases struggles to scale with the complexity of modern multi-omics data (genomic, transcriptomic, proteomic), leading to incomplete or outdated target identification (Paananen & Fortino, 2020; Zhou et al., 2022; Trajanoska et al., 2023; Lindsay, 2003; Zhou & Zhong, 2017). Second, traditional network-based prioritization, which prioritize genes based on protein-protein interaction (PPI) network centrality, oversimplify biological context by ignoring tissue-specific genomic features such as mutation frequencies and differential expression profiles (Petti et al., 2020). Third, fragmented prioritization—reliance on single-metric approaches like fold change or mutational frequency—introduces variability due to arbitrary thresholds and sample bias (McCarthy & Smyth, 2009; Dinstag & Shamir, 2020; López-Cortés et al., 2018).

These gaps contribute to high failure rates in translating preclinical discoveries to clinical therapies, particularly in genetically heterogeneous cancers like pancreatic cancer (Singh et al., 2023; Sun et al., 2022; Zhu et al., 2021; Somarelli et al., 2019).

Computational advances address these challenges by integrating multi-omics data, network-based prioritization and AI-driven literature review, driving down costs and expediting the development of effective therapies through in silico assessments (Sadybekov & Katritch, 2023; Sliwoski et al., 2014; Huan et al., 2010). These approaches harness the power of neural networks, expansive genomic datasets, and sophisticated machine learning algorithms to predict genes of interest with a precision unattainable by traditional methods (Chen et al., 2006). The integration of multi-omics data contextualizes mutations within tissue-specific expression patterns, while network-based prioritization refines prioritization by mapping genes to functionally relevant pathways (Shim et al., 2015). Concurrently, AI-driven literature review (e.g., GPT-4) automates the synthesis of preclinical and clinical evidence, identifying targets with mechanistic and translational relevance (Liu et al., 2021; Oniani et al., 2024; Sallam, 2023; Tripathi et al., 2024). Together, these components resolve biases inherent to fragmented analyses: multi-omics data anchors targets in biological context, networks prioritize functional relevance, and AI-driven literature review ensures scalability in evidence synthesis. (Somarelli et al., 2019; Zhu et al., 2021; Sadybekov & Katritch, 2023).

Multi-Omics data integrates genes based on genomic information (Eg. Differential Gene Expression and Frequency of Mutation). Differential gene expression is a critical method for identifying genes significantly altered between conditions, such as cancerous versus normal tissues (Bai et al., 2013; Van de Sande et al., 2023). A common approach involves calculating "fold change," which quantifies the ratio of gene expression levels between these states (Love et al., 2014; Mutch et al., 2002). However, the arbitrary selection of fold change thresholds can introduce variability into prioritization, potentially compromising the reliability of target identification (McCarthy & Smyth, 2009). Tools like GEO2R exemplify this approach, leveraging fold change to rank genes under experimental conditions, such as tumor versus healthy tissue comparisons (Barrett et al., 2013). Separately, frequency-based prioritization methods focus on genes with elevated mutational rates in disease contexts, hypothesizing these as common therapeutic targets (Dinstag & Shamir, 2020; López-Cortés et al., 2018). While valuable, frequency-based prioritization methods for gene prioritization can be prone to bias, especially due to sample selection, which can skew results (Lazzeroni et al., 2014). To address these limitations, network centrality-based prioritization has emerged as a complementary strategy. This approach leverages gene connectivity within biological networks, offering a holistic framework for target selection by expanding gene lists and strengthening disease association metrics (Janyasupab et al., 2021; Magger et al., 2012).

Network-based prioritization enables researchers to analyze genomic datasets and identify critical regulatory genes implicated in cancer development (Chang et al., 2021; Sonehara & Okada, 2021). These methods prioritize disease-related genes by integrating data from PPI networks and known gene-drug associations (Mohsen et al., 2021; Zhang et al., 2021). The ability to efficiently process genomic information and derive meaningful insights is pivotal for identifying and visualizing relevant drug targets, underscoring the importance of network-based prioritization (Chen et al., 2013; Chen et al., 2009; Huan et al., 2010). Network-based prioritization methods are particularly favored for gene prioritization, as they utilize established biological networks to identify genes associated with specific diseases (Shim et al., 2015; Huang et al., 2012). As noted earlier, traditional network-based prioritization methods lack tissue specificity (Petti et al., 2020), necessitating the integration of genomic data.

As noted earlier, AI-Driven Literature review through the use of Large Language Models (LLMs), such as GPT-4, have emerged as another transformative *in silico* approach to drug prioritization. ( Liu et al., 2021; Oniani et al., 2024). LLMs can predict essential information about gene targets, including structural domains of proteins, protein structure, toxicity and adverse effects, functional significance, clinical and preclinical relevance, and treatment efficacy (Sallam, 2023; Tripathi et al., 2024). Furthermore, GPT-4 has demonstrated the ability to rival human performance in conducting literature reviews, thus streamlining a critical component of the drug target prioritization process (Khraisha et al., 2024; Li et al., 2010)

In this study, we hypothesize that the utilization of network-based analysis, artificial intelligence, and biologically significant data will enable systemic prioritization of actionable therapeutic targets. Thus, we propose GETgene-AI, a framework which annotates network-based analysis with LLM enabled literature review, and biologically significant data. Central to GETgene-AI is the G.E.T. strategy, which integrates three key data streams: the G List (genes with genetic mutations, variations functionally implicated in genotype-to-phenotype association studies of the disease), the E List (disease target tissue-specific expressions of the candidate gene), and the T List (established drug targets based on reports from literature, patents, clinical trials, or existing approved drugs). Initial gene candidates are derived from heterogeneous biological datasets; including fold change, copy number alterations, and mutational frequency metrics. To mitigate biases inherent to fragmented or incomplete data, GETgene-AI employs a multi-dataset integration approach. The framework iteratively refines candidate lists through the network-based tool BEERE, which annotates and prioritizes genes with network-based centrality methods to create a high-quality, prioritized gene list. This iterative process expands and ranks candidates by evaluating their biological relevance, network centrality, and concordance with genomic aberrations, thereby improving target identification accuracy. GPT-4o is integrated into the process to improve literature review efficiency and further annotate the target list, enhancing the overall workflow. By combining traditional and *in silico* methods, GETgene-AI bridges gaps in drug discovery and facilitates the development of personalized cancer therapies.

The novel drug targets prioritized through our case study in pancreatic cancer not only offer insights into the unique molecular mechanisms driving this aggressive cancer but also present promising avenues for therapeutic intervention. While pancreatic cancer serves as a case study in this paper, the underlying methodology is adaptable to a wide range of cancers and diseases, thereby accelerating the discovery of therapeutic options.

## METHODS

In figure 1, we show a general overview of the GETgene-AI framework.

**Figure 1:**
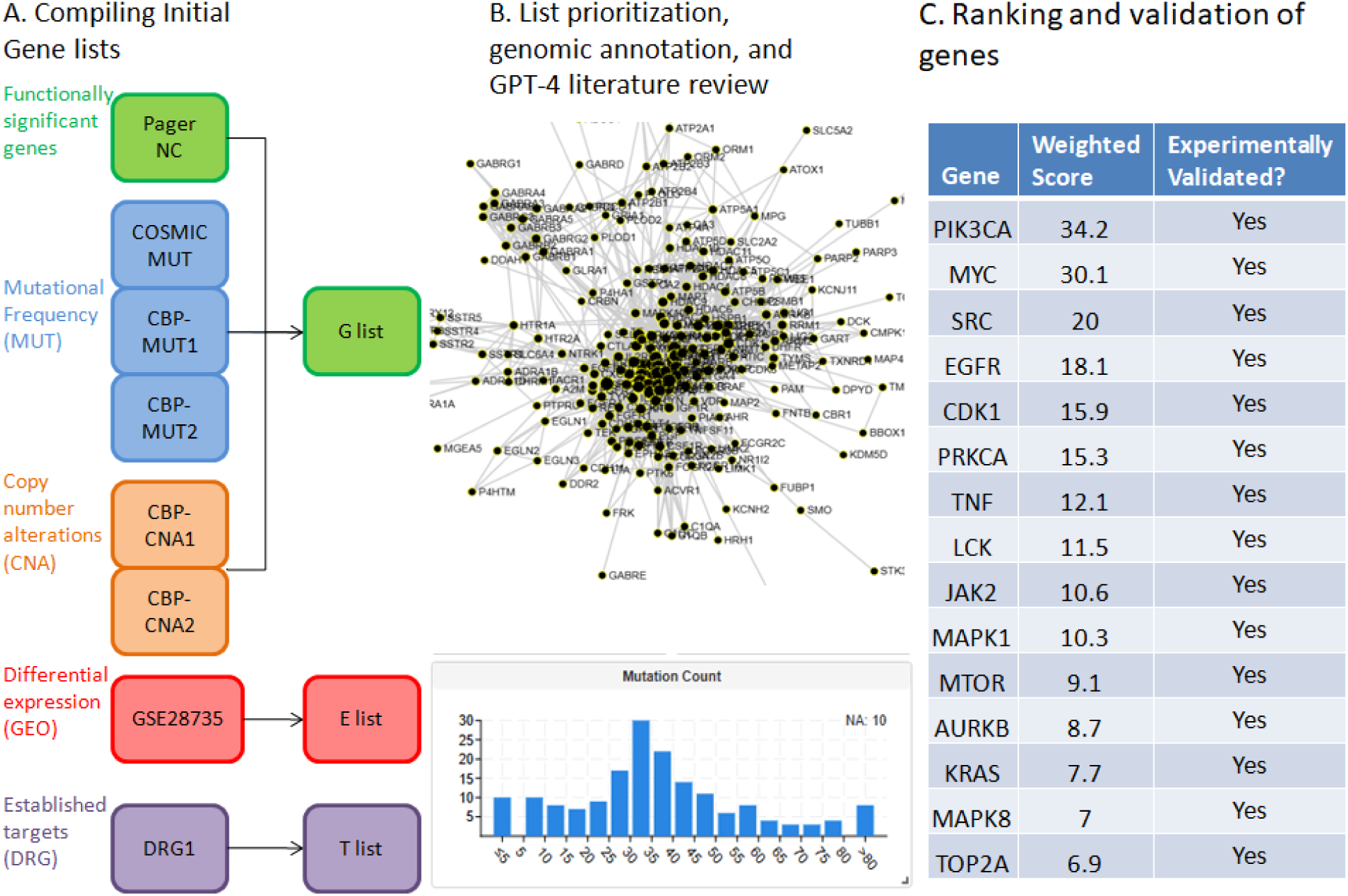
General overview of the GET list compilation and ranking process. A. Initial gene lists from each of the three subsets are compiled. 2493 genes are compiled in the initial G list, 2000 genes are compiled in the initial E list, and 131 genes are compiled in the initial T list. B. Each list is prioritized using the BEERE network ranking and expansion tool, before being annotated with genomic information and GPT4 literature review. C. A weighted score is calculated to rank the list and genes are manually validated through literature review.

The initial gene list is generated by employing a three-tiered strategy—comprising the Gene list (G list), Expression list (E list), and Target list (T list)—to integrate biological context into gene prioritization. The G list identifies genes with high mutational frequency, functional significance (e.g., pathway enrichment via the Kyoto Encyclopedia of Genes and Genomes (KEGG)), and genotype-phenotype associations. The E list focuses on genes exhibiting significant differential expression in pancreatic ductal adenocarcinoma (PDAC) compared to normal tissues, while the T list incorporates genes annotated as drug targets in clinical trials, patents, or approved therapies. To construct these lists, disease-specific genomic data were aggregated from public databases (e.g., TCGA, COSMIC, PAGER) and processed using GRIPPs (Gong & Chen, 2023), an iterative network-based approach that applies modality-specific thresholds to ensure robust inclusion criteria.

Following the initial gene list generation, the second step involves prioritizing and expanding these lists using the BEERE network-ranking tool. BEERE was selected for its demonstrated efficacy in filtering low-confidence data and enhancing prioritization accuracy(Yue et al., 2019), ensuring comprehensive and reliable gene sets.

In the third step, genes implicated in clinical trials were incorporated and annotated with factors relevant to drug target prioritization. These included factors such as clinical trial popularity, differential expression, mutation frequency, and copy number alterations. This process is further enriched by leveraging GPT-4 enabled literature review, which provides additional context and insights into the biological and clinical relevance of the identified genes.

Finally, the GETgene-AI ranking is generated by creating a weighted score called RP-score which integrates genomic information, BEERE list scores, and expression in important organs to assess the efficacy of targeting a gene. The weights for each variable were determined by calculating Spearman correlations between the specific variable and clinical trial popularity, defined as the number of cancer treatment–related clinical trials in which the gene was targeted.

Using PDAC as a case study (selected due to its poor prognosis and limited therapeutic options ( Hu et al., 2021)) our framework produced quantitative data and novel insights into potential therapeutic targets, demonstrating its utility in advancing precision oncology.

### Initial gene list generation

### 1) Compiling the Gene list from Genetic Mutations

For the "GENE" component of our "GET" framework, we compiled three gene subsets: PAGER-NC, COSMIC-MUT, and CBP-CNA-MUT. The initial "GENE" list was compiled from the PAGER ( Huang et al., 2012; Yue et al., 2018, 2022), cBioPortal (de Bruijn et al., 2023), and COSMIC (Tate et al., 2019) databases. To address potential sample biases and data incompleteness (e.g., studies failing to detect specific genes), we incorporated multiple datasets from these repositories when available. Genes associated with the term "Pancreatic Cancer" were manually curated from these databases. Empirical cutoffs were applied to prioritize genes with relevance to pancreatic cancer.

To integrate biological pathway context into gene prioritization, we utilized PAGER (Chowbina et al., 2009), which quantifies functional significance through pathway-based metrics. From PAGER, 844 candidate genes were selected heuristically using an nCoCo score threshold between 5 and 100. We integrated nCoCo score since it measures coherence by integrating co-citation and pathway data, with higher scores indicating stronger biological cohesion ( Huang et al., 2012; Yue et al., 2018, 2022). Groups with nCoCo scores below 5 lacked coherent pathway annotations or shared functional roles (e.g., random co-citation clusters), reducing their relevance to disease mechanisms. Groups with nCoCO scores above 100 represented ubiquitous biological processes (e.g., "cellular metabolism") with broad roles across tissues, diluting disease-specific signal.

For the cBioPortal and COSMIC databases, thresholds were defined by identifying points where mutational frequency no longer demonstrated cancer-specific significance in prior studies. From cBioPortal, 1,000 genes were selected using cutoffs of 8.2% for copy number alterations (CNA) and 2.8% for mutational frequency. The threshold for copy number alterations is significantly higher due to only 21 sets of copy number signitures being represented in 97% of tumor samples on The Cancer Genome Atlas (Steele et al., 2022). The 2.8% cutoff for mutational frequency is due to the fact that a limited amount of genes were found to be mutated in more than 5% of tumors (Sinkala, 2023). Most biologically relevant genes were found to be mutated at frequencies between 2-20% (Lawrence et al., 2014). From COSMIC, 649 genes were compiled using a 20% mutational frequency cutoff according to the previously mentioned frequency range. To prevent data bias, datasets with differing mutational frequency within samples of the same genes were selected. Integration of multiple datasets was also utilized to prevent sample bias. Finally, candidate genes from PAGER, cBioPortal, and COSMIC were aggregated to form the “G list”, comprising 2,493 genes in total.

While we did not formally reprocess the entire dataset with varied cutoffs, preliminary assessments suggested that slight variations (e.g., ±5% in mutation frequency thresholds) did not significantly alter the prioritized gene rankings. This stability indicates that GETgene-AI’s prioritization is driven by biologically relevant signals rather than arbitrary thresholds.

### 2) Compiling candidate genes for the “Expression” subset

Candidate genes were prioritized by analyzing the GEO dataset GSE29735, titled "Pancreatic ductal adenocarcinoma tumor and adjacent non-tumor tissue" (Zhang et al., 2012, 2013), using the GEO2R tool. Samples were categorized into tumor and non-tumor groups via the "Define groups" feature, with the tumor group defined as "human pancreatic tumor tissue patient samples" and the non-tumor group as "human pancreatic non-tumor patient samples". The dataset comprised of 90 patient samples, evenly distributed between 45 tumor and 45 non-tumor samples. Differentially gene expression analysis was performed using GEO2R’s "analyze" function. The top 2,504 genes exhibiting logfc values over 0.25 were compiled into an initial “E list”. A cutoff of 0.25 was determined based on the “FindAllMarker” function provided by the R package Seurat (Wang et al., 2024). The list was subsequently processed iteratively using the BEERE software in accordance with the GRIPPs method.

### 3) Compiling candidate genes for the “Target” subset

Incorporating pharmacology data with network-based prioritization is a well established approach( Huang et al., 2015; Huang et al., 2012b). Building on this methodology, a set of 131 genes were identified using DrugBank (Wishart et al., 2018), a comprehensive drug and drug-target database. To extract relevant genes, the database was queried using the search terms “Pancreatic Cancer,” “Pancreatic Ductal Adenocarcinoma,” and “Neuroendocrine Pancreatic Cancer” within its drug repository. Drugs explicitly indicated for Pancreatic Cancer treatment were identified by reviewing their associated metadata, including summaries, background descriptions, indications, clinical trial references, and listed “Associated Conditions.” Each drug’s mechanism of action, therapeutic summary, and clinical trial references were manually evaluated to distinguish agents directly treating pancreatic cancer from those used for supportive care (e.g., chemotherapy relief, pain management, or sedation). For all drugs meeting the inclusion criteria, gene targets listed under their respective “Targets” section in DrugBank were compiled, resulting in 131 unique genes associated with pancreatic cancer therapeutics.

### Prioritization and expansion of GET lists

To improve the specificity and biological relevance of our candidate gene lists, we implemented an iterative refinement process using the BEERE tool for prioritization and network-based expansion. The BEERE tool employs an initial ranking algorithm and two iterative ranking algorithms—PageRank and an ant-colony algorithm—both of which have demonstrated success across diverse knowledge domains (Yue et al., 2019). Although both ranking algorithms use an iterative ranking process, they differ in how node importance weights are calculated. The PageRank algorithm assigns node importance directly from neighboring nodes. In the ant-colony algorithm nodes lose score when disseminating information and gain score upon receiving it. BEERE expands the gene list using the nearest-neighbor network constructed from protein-protein interactions in the HAPPI 2.0 database (Chen et al., 2009, 2017; Wu et al., 2012).

This workflow addresses the inherent limitations of single-dimensional analyses (e.g., relying solely on mutation or expression data) by integrating complementary biological evidence. Building on the GRIPPS framework (Gong & Chen, 2023), we developed a customized pipeline to systemically prioritize genes from three distinct categories: the combined GET list(genes ranked by aggregated mutational frequency, differential expression, and known drug-target status), the GT list (genes co-occurring in mutation and drug-target databases to highlight functionally relevant drivers,) and a prioritized Expression (E) list (genes ranked exclusively by differential expression in pancreatic cancer) . This systematic approach ensured a robust and multifaceted analysis of candidate genes, enhancing the identification of potential therapeutic targets.

Each list underwent the same refinement workflow to balance comprehensiveness with specificity. First, BEERE expanded the initial gene sets by incorporating proximal interactors from protein-protein interaction (PPI) networks in the HAPPI 2.0 database, thereby capturing functionally related genes beyond those directly identified in our initial screens. Next, BEERE’s network propagation and statistical ranking algorithms prioritized genes based on their network centrality and significance scores. To prevent overexpansion and maintain focus on high-confidence candidates, we empirically filtered each list to retain the top 500 genes after each prioritization cycle. This iterative process was repeated three times, as preliminary testing revealed that additional iterations caused excessive convergence of the lists, reducing their distinct biological relevance. Three iterations optimally preserved the unique profiles of each list while still enabling meaningful integration.

The refined outputs were consolidated into an Initial GET List, which underwent a final BEERE-based prioritization to generate the Final GET List. For comparative analysis, we also derived two additional lists: the GT List (prioritized genes from mutation-drug target overlaps) and the Expression List (top differentially expressed genes). This tiered approach ensured that our final candidate pool retained both mechanistic diversity (genes linked to distinct biological processes) and clinical relevance (genes with actionable potential as drug targets).

This refinement process was critical to address three key challenges: (1) mitigating the high false-positive rate inherent to mutational and expression screens in heterogeneous cancers like pancreatic adenocarcinoma, (2) reconciling discrepancies between genes prioritized by individual data types (e.g., highly mutated genes often lack expression changes, and vice versa), and (3) ensuring functional coherence by embedding candidates within PPI networks reflective of disease biology. By iteratively refining lists through network propagation and multi-evidence integration, we enhanced the biological plausibility of candidates while preserving distinct mechanistic hypotheses for downstream validation.

### GPT-4o aided literature assessment

Recent research has demonstrated that GPT-4o performs "human-like" literature reviews, particularly in screening and analyzing scientific literature (Khraisha et al., 2024). For this study, abstracts related to pancreatic cancer genes and treatments were downloaded using PubMed’s "save" feature. A total of 5,091 abstracts were collected and uploaded for analysis by GPT-4o through a custom GPTo interface. Due to the data processing limitations of GPT-4o, abstracts were filtered to include only meta-analyses, clinical trials, and systematic reviews on PubMed to ensure high-quality input data.

GPT-4o was configured with specific instructions to rank genes based on a scoring system with a maximum score of 400 points, distributed across four categories: functional significance in pancreatic cancer, research popularity, treatment effectiveness when targeting or inhibiting the gene, and protein structure. Each category was allocated 100 points, and the resulting metric was termed the GPT-4 score.

To mitigate GPT-4o’s known issue of "hallucination" or the generation of inaccurate or nonexistent information, the model was explicitly instructed to base its rankings solely on the uploaded research database. Additionally, the model was required to cite articles referenced during the ranking process and provide explanations for the scores assigned to each gene in every category. GPT-4o outputs were manually verified against curated datasets to ensure biological relevance and mitigate hallucinations. Citations provided by GPT-4o were cross-referenced with PubMed to confirm validity. All cited articles were manually verified, and any errors or hallucinations were addressed by instructing the model to re-search the uploaded literature database for accurate mentions of the gene.

### Incorporation of clinically implicated genes and annotation of genes with factors relevant to drug target prioritization

Clinical trials are critical for evaluating the efficacy of therapeutic agents targeting specific genes. To assess the clinical relevance of prioritized genes, we quantified clinical trial activity by compiling the frequency of trials associated with each gene. Genes targeted by drugs investigated in pancreatic cancer treatment trials systemically identified through the following process: A search for the term “pancreatic cancer” was conducted on Clinicaltrials.gov, and all drugs listed in active or completed interventional trials for pancreatic cancer were extracted.

Corresponding target genes for these drugs were then identified using DrugBank’s “Targets” section, which provides genes targeted by the drug for pancreatic cancer treatment. This process yielded 357 drugs targeting 253 unique genes. These genes were annotated with BEERE scores derived from the previously described GET lists. To enhance biological validity, the analysis integrated quantitative genomic datasets. Mutation frequency data was obtained from cBioPortal (de Bruijn et al., 2023), while protein expression profiles across tissues relevant to therapeutic safety (e.g, brain, gastrointestinal tract, liver, and kidney) were sourced from the ProteinAtlas (Uhlén et al., 2015).

Following the prioritization of the GET list and identification of clinically trialed genes, we annotated these genes with functional genomic data. Mutational frequency—a key determinant in gene ontology ranking (Timar & Kashofer, 2020)—and Copy Number Alterations **(CNA)**, a critical marker of genomic instability (Beroukhim et al., 2010), were evaluated .

Mutation and CNA data were sourced from CBioPortal (de Bruijn et al., 2023) using two cohorts: the “Pancreatic Cancer (UTSW, Nat Commun 2015)” and “Pancreatic Adenocarcinoma (TCGA, PanCancer Atlas)” studies, both of which employed whole-exome sequencing for all samples.

Tissue-specific expression is a vital factor in gene prioritization (Beroukhim et al., 2010). Genes with high expression in essential tissues—such as the heart, liver, gastrointestinal system, brain, and kidneys—pose a higher risk of adverse effects when targeted, necessitating their de- prioritization. Annotation of tissue expression was performed using the “RNA expression score” provided by ProteinAtlas (Uhlén et al., 2015), a comprehensive database mapping protein expression in various organs. This RNA expression score, manually calculated, measures the RNA expression levels of genes across different tissues.

### GETgene-AI ranking

To unify these criteria, we developed a weighted RP score that integrates mutation frequency, CNA, mutational frequency, GET list scores, tissue expression, E list scores, GT list scores, and clinical trial popularity (number of trials testing drugs targeting a given gene for cancer therapy). Spearman correlation analysis was performed to assess pair wise relationships between these variables. The Spearman correlation is a widely used statistical tool for prioritizing drug target discovery, as it quantifies the strength and direction of monotonic associations between variables (Tanoli et al., 2025). Unlike the Pearson correlation, which measures linear proportional relationships, Spearman’s method evaluates rank-based correlations, making it less sensitive to outliers—a critical advantage when analyzing noisy biological datasets (Rodríguez-Pérez & Bajorath, 2021). Additionally, Spearman correlation captures both linear and nonlinear monotonic relationships, enabling researchers to identify diverse patterns in genomic data that might otherwise be overlooked (Penrod & Moore, 2014).

These features collectively make it a versatile choice for exploring complex biological associations in drug target discovery pipelines. The RP score weights were derived from the directional Spearman correlation coefficients (ρ) between clinical trial popularity and each genomic/contextual variable. Unlike absolute correlations, this approach preserves the sign of the relationship, allowing variables with positive associations (e.g., higher mutation frequency → more trials) to enhance the RP score, while variables with negative associations (e.g., higher tissue expression → fewer trials) reduce it. This directional weighting reflects the observed biological and clinical trends, where certain genomic features either promote or deter therapeutic targeting. The RP score provides a multi-dimensional ranking mechanism for therapeutic target prioritization. GTlistscore received the highest weight because it integrates established drug targets and functionally or frequently mutated genes into its score. Table 1 summarizes the relative weights of each factor in the RP score, ranked in descending order of contribution.

**Table 1:**
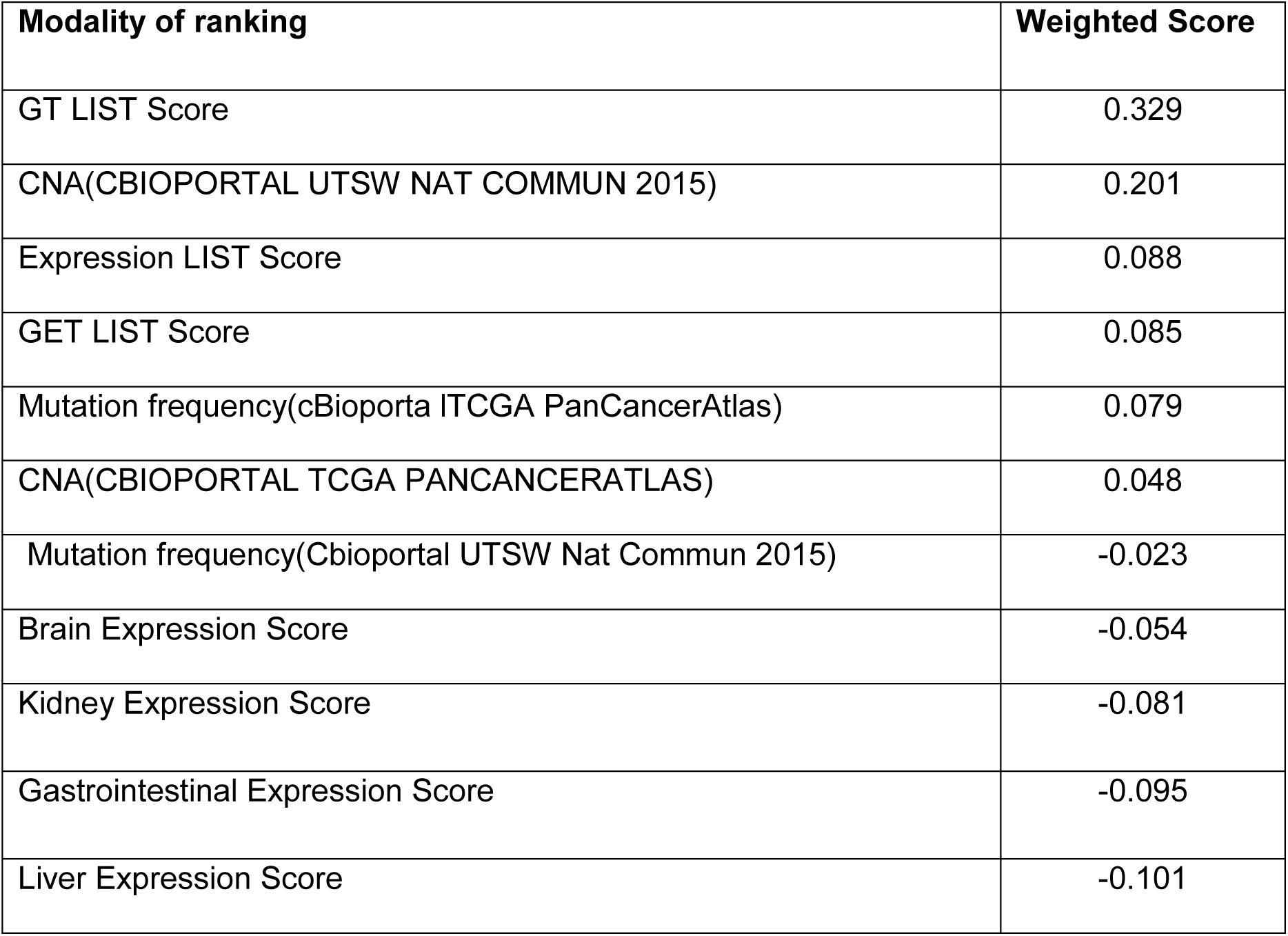
Weights each modality was assigned for calculation of the RP score in GETGENE-AI.

### Mitigation of Bias and False Positives

To address potential sample biases and data incompleteness—such as studies failing to detect specific genes—multiple datasets from the same databases were utilized wherever possible. This redundancy ensured a more comprehensive analysis and minimized the impact of dataset-specific variability. For example, multiple studies within CBioPortal, such as “Pancreatic Cancer (UTSW, Nat Commun 2015)” and “Pancreatic Adenocarcinoma (TCGA, PanCancer Atlas),” were analyzed concurrently to increase the reliability of mutational frequency and CNA data. We combined data sources after thresholding as a form of preprocessing to ensure stable rankings, and reducing sensitivity to arbitrary thresholds. Mutational and CNA data were merged from multiple studies to reduce cohort specific bias.

While we did not formally reprocess the entire dataset with varied cutoffs, preliminary assessments suggested that slight variations (e.g., ±5% in mutation frequency thresholds) did not significantly alter the prioritized gene rankings. This stability indicates that GETgene-AI’s prioritization is driven by biologically relevant signals rather than arbitrary thresholding.

To further enhance the accuracy of the prioritization process, each gene within the top 250 ranked by RP score was manually verified through a literature review to confirm its role in cancer biology. This step was critical in identifying and eliminating false positives. Notably, no genes within the top 250 were found to be false positives, validating the robustness of the RP scoring methodology.

Additionally, hallucination errors from GPT-4o were mitigated through a structured training approach. The model was instructed to explicitly cite a source used in the calculation of each gene’s ranking score. These citations were manually evaluated for accuracy and relevance, ensuring that the ranking process was grounded in verifiable scientific evidence. This dual-layered validation—automated scoring combined with manual review—was integral to maintaining the integrity and reliability of the gene prioritization framework.

### Statistical Methods

Spearman correlation coefficients were computed to assess the alignment of GPT-4o rankings with network-derived rankings. The Spearman correlation between the GPT-4 score and the Weighted Score was 0.291, indicating some significance. Interestingly, GPT-4 score is more strongly correlated with all BEERE list ranking scores, with 0.478 between GPT-4 score and Expression list score, 0.457 between GPT-4 score and Combined weighted score of all BEERE lists, 0.454 correlations between GPT-4 score and GET list score, and 0.444 between GPT-4 score and GT list score. These results indicate that the GPT-4 score is more similar to that of standard network prioritization techniques, which may be a result of the training data utilized.

### Comparing research relevance to rank on GETgene-AI

To compare the popularity to the rankings of each gene in both the GPT-4 Score and the RP scores, the amount of results contained on PubMed when searching “Gene name Pancreatic Cancer” were compiled and used for the GPT-LIT score, and the RP-LIT score. The GPT-LIT score is the GPT4-score divided by the amount of publications on PubMed, while the RP-lit score is the RP-score divided by the amount of publications on PubMed. Genes with no functional relationship to cancer in any way were excluded from the rankings to remove false positives.

## RESULTS

### GETgene-AI Rankings and Validations

The highest ranked genes according to GETgene-AI in table 2.

**Table 2:**
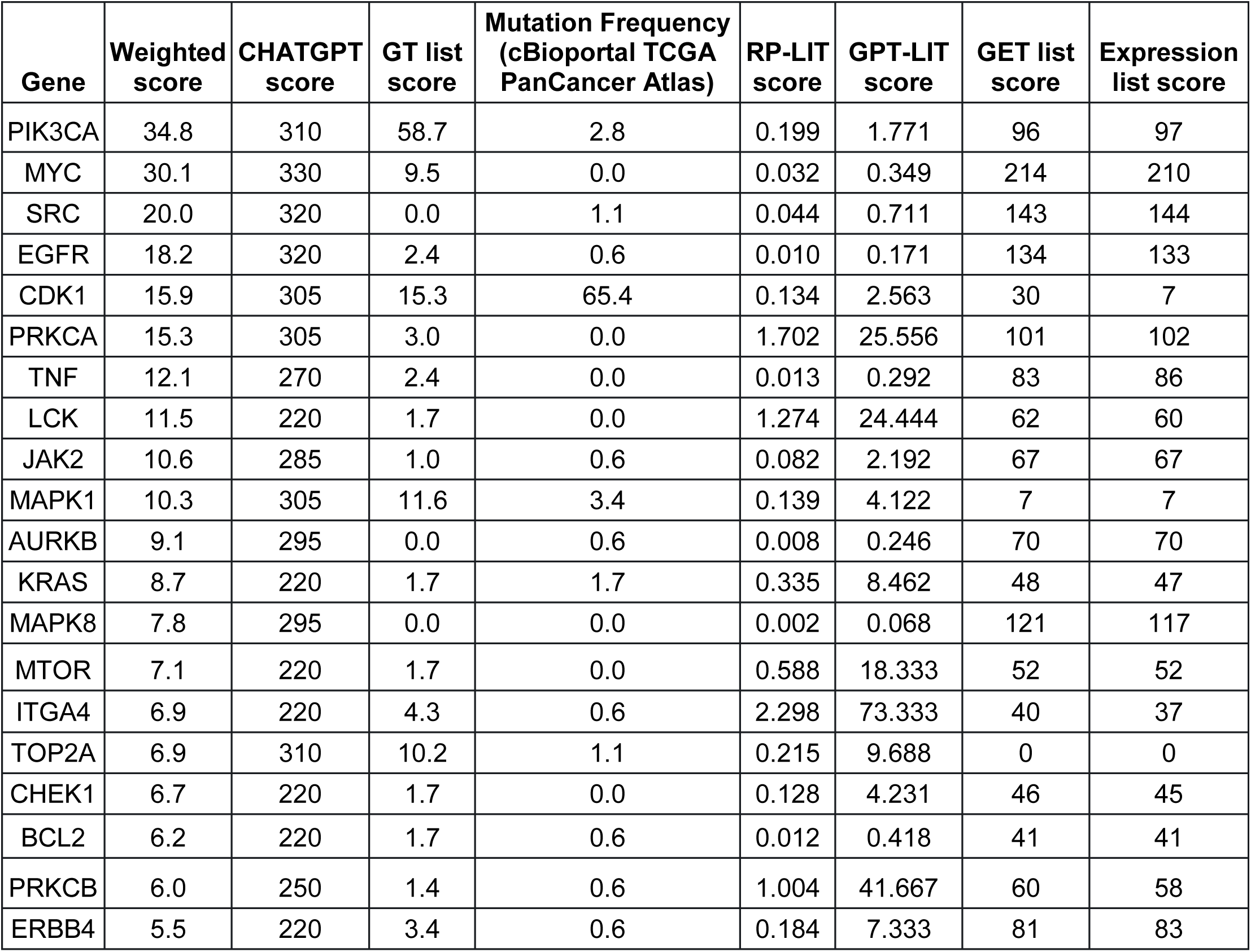
Highest 20 genes ranked on GETGENE-AI. Weighted score is RP score, CHAT GPT score is GPT4 score.

*PIK3CA* emerged as the highest-ranked gene on our list. It encodes the enzyme PI3K, which regulates critical cellular processes such as growth, metabolism, proliferation, and apoptosis(Conway et al., 2019). *PIK3CA* also modulates downstream effectors, including *AKT* and *mTOR* (Ala, 2022), and preclinical studies demonstrate that mutations in this gene sensitize cancers to dual *PI3K*/*mTOR* inhibitors (Zhang et al., 2021), underscoring its therapeutic potential. Notably, *PIK3CA*-null tumors exhibit heightened susceptibility to T-cell surveillance *in vitro* (Sivaram et al., 2019), while its inhibition in pancreatic cancer models initiates tumorigenesis (Payne et al., 2015), highlighting its dual role in progression and therapy.

*MYC*, the second highest-ranked gene, achieved its position due to its top GET list score, reflecting its network centrality among the 500 most expressed, clinically relevant, and frequently mutated genes. Overexpression of *c-MYC* is a hallmark of aggressive pancreatic cancer, where it binds promoter regions of oncogenic targets (Hayashi et al., 2021). Despite its pivotal regulatory role, *MYC*’s complex protein structure poses therapeutic challenges, resulting in a lower GT list score. Recent advances in small-molecule inhibitors, however, show preclinical promise.

*SRC* ranks as the third-highest gene on our list, driven by its high scores in both the GET list and Expression list modalities. Inhibition of *SRC* in pancreatic cancer has been shown to reverse chemoresistance to pyroptosis in both in vitro and in vivo studies (Su et al., 2023).

Aberrant *SRC* activity promotes tumorigenesis and is frequently associated with poor prognosis in pancreatic ductal adenocarcinoma (PDAC) (Poh & Ernst, 2023). Several *SRC*-targeting therapies are currently under clinical investigation (Hilbig, 2008).

*EGFR* is the fourth highest-ranked gene, attributed to its high GET list and Expression list scores. *EGFR* is not ranked higher due to having lower mutational and copy number alteration frequencies than other high ranking targets. *EGFR* is also implicated in tumorigenesis, particularly in lung and breast cancer (Sigismund et al., 2018). Anti-*EGFR* agents have shown significant clinical promise, despite associated adverse effects (Verma et al., 2020).

*KRAS* ranks twelfth on our list, despite its prominence in pancreatic cancer research, with over 4,545 PubMed articles on *KRAS* mutations in pancreatic cancer. However, it is ranked lower due to its low differential expression, being ranked 27562 on GEO2R. Additionally, KRAS is considered undruggable due to the lack of classic drug binding sites, further increasing difficulty in targeting it (Zhu et al., 2022). The KRAS oncogene plays a critical role in the initiation and maintenance of pancreatic tumors (Luo, 2021). *KRAS* mutations are present in over 90% of PDAC cases, but therapeutic inhibition remains highly challenging, with effective inhibitors only recently being discovered (Bannoura et al., 2021).

*CDK1* ranks fifth on our list, largely due to its high scores in both the GET and Expression lists. *CDK1* is strongly correlated with prognosis and is highly expressed in pancreatic cancer tissue, as well as in response to gemcitabine, an approved pancreatic cancer drug (Xu et al., 2023). Additionally, inhibition of *CDK1*, along with *CDK2* and *CDK5*, has been shown to overcome IFN-γ-triggered acquired resistance in pancreatic tumor immunity (Huang et al., 2021).

*PRKCA* ranks seventh on our list. It encodes protein kinase C and is mutated in various cancers. *PRKCA*’s high ranking is attributed to its strong GET and Expression list scores, as well as its extremely low organ expression score. It is strongly associated with the activation of the protein translation initiation pathway (Rosenberg et al., 2018) and is a hallmark mutation in chordoid gliomas (Jiang et al., 2019). *PRKCA* also contributes to susceptibility to pancreatic cancer through the peroxisome proliferator-activated receptor (*PPAR*) signaling pathway, which plays a key role in pancreatic cancer development and progression ( Liu et al., 2020). Inhibition of *PRKCA* has demonstrated antitumor activity in patients with advanced non-small cell lung cancer (NSCLC) (Villalona-Calero et al., 2004).

*TNF* is the eighth highest-ranked gene on our list. Tumor Necrosis Factor (*TNF*) upregulation is associated with invasion and immunomodulation in pancreatic cancer (Wiedmann et al., 2023). *TNF*-mutated macrophages have also been shown to promote aggressive cancer behaviors through lineage reprogramming (Tu et al., 2021).

*LCK* ranks ninth on our list. This gene is expressed in tumor cells and plays a key role in T-cell development (Bommhardt et al., 2019). High *LCK* protein expression has been associated with improved patient survival in cancer (Cancer Genome Atlas Network, 2015). Despite its biological relevance, *LCK* has only four PubMed publications discussing its role in pancreatic cancer as of May 2024. Its identification as a high-priority target demonstrates GETgene-AI’s ability to prioritize genes with strong biological relevance but limited literature prominence.

*ITGA4* ranks 15th on our list. It has an extremely low organ expression score and only four PubMed articles discussing its role in pancreatic cancer. *ITGA4* has potential as an independent prognostic indicator for patient survival and has been linked to the *PI3K*/*AKT* pathway (Faleiro et al., 2021). Its identification as a high-priority target further highlights GETgene-AI’s capability to prioritize genes with strong biological relevance despite limited literature attention.

*KCNA* ranks 34th on our list. Notably, there are no PubMed publications describing its relation to pancreatic cancer, and only three publications mention its role in cancer in general. The identification of *KCNA* as a high-priority target underscores GETgene-AI’s ability to prioritize genes with strong biological relevance but minimal literature prominence. *KCNA* exhibits differentially high expression in stomach and lung cancers and is positively correlated with infiltrated immune cells and survival rates (Angi et al., 2023).

During the iterative ranking process, genes lacking functional relevance to cancer were systematically deprioritized. For instance, genes that ranked highly due to algorithmic artifacts but lacked experimental validation or literature support were ranked lower than genes with experimental validation or literature support.

### Comparing GETgene-AI to other frameworks

We benchmarked GETgene-AI against two other frameworks: one focused on differential expression analysis and the other on network-based gene prioritization. We chose to benchmark off of precision, defined as the percentage of genes found to be functionally significant in literature. PubMed was utilized to search for literature in this benchmark. For the differential expression comparison, we selected GEO2R, utilizing the GSE28735 dataset, which was integrated into the ’Expression list’ component of our GET lists. Genes were ranked based on their log-fold change (log-fc), representing the difference in gene expression between tumor and non-tumor groups. In the GEO2R list, the top-ranked genes were *PNLIPRP1* and *PNLIPRP2*, both of which encode pancreatic lipase-related proteins critical for digestion and fat absorption (Zhu et al., 2021). However, these genes are not considered viable targets for pancreatic cancer. The third-ranked gene, *IAPP* (Islet Amyloid Polypeptide), has been shown to lack tumor suppressor functionality, and loss of *IAPP* signaling is not associated with pancreatic cancer (Taylor et al., 2023). Among the top 50 genes identified by GEO2R, 30 were experimentally validated as relevant to pancreatic cancer. In contrast, GETgene-AI prioritized 49 experimentally validated targets within its top 50, representing a 38% improvement over GEO2R. GEO2R’s limitations, including the absence of mutational frequency analysis, functional impact assessment, network-based analysis, and adverse effect evaluation, hinder its utility in drug target discovery. In comparison, GETgene-AI leverages statistical filtering and incorporates genomic information, significantly enhancing both the efficiency and quality of gene prioritization. Figure 2 presents a volcano plot illustrating the log2(fold change) distributions for the analyzed genes.

**Figure 2:**
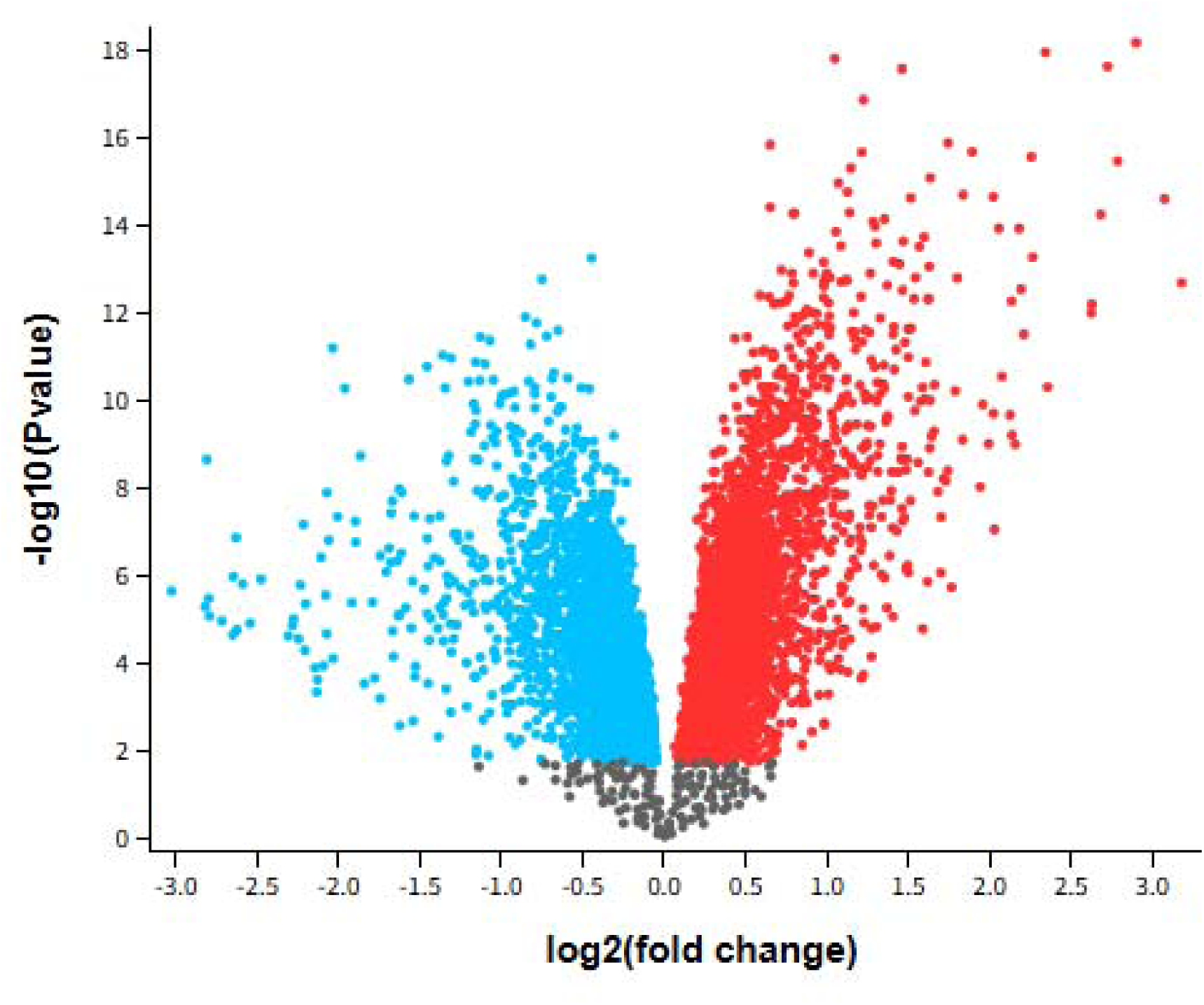
Volcano plot GSE28735: Microarray gene-expression profiles of 45 matching pairs of tumor vs. nontumor, Padj<0.05. Blue indicates down regulated while red indicates upregulated.

For the network-based comparison, we employed STRING, a database that integrates protein-protein interaction data (Szklarczyk et al., 2023), focusing specifically on the KEGG pathway hsa0512 (Kanehisa, 2019; Kanehisa et al., 2025; Kanehisa & Goto, 2000). Genes were ranked based on node degree, a measure of the number of interactions a protein has within the network (Bozhilova et al., 2019). The highest-ranked gene in the STRING list was *AKT1*, a protein kinase known to stimulate cell growth and proliferation (Grassilli et al., 2020).

However, *AKT1* has been shown to resist inhibition by shifting its metabolic activity from glycolysis to mitochondrial respiration (Arasanz et al., 2019). Additionally, it exhibits a low mutational frequency of only 1% in a cohort of 19,784 patients with various tumors (Millis et al., 2016). Due to its low mutational frequency and the challenges associated with its inhibition, *AKT1* was ranked 33rd by GETgene-AI. Among the top 50 genes prioritized by STRING, 47 were experimentally validated for relevance to pancreatic cancer, whereas GETgene-AI identified 49 experimentally validated genes within its top 50, demonstrating an improvement over STRING. STRING’s limitations, such as its inability to account for mutational frequency and other critical factors in drug target identification, result in a narrower focus, with only 81 targets prioritized compared to the more comprehensive analysis provided by GETgene-AI. Figure 3 illustrates the network constructed using STRING.

**Figure 3:**
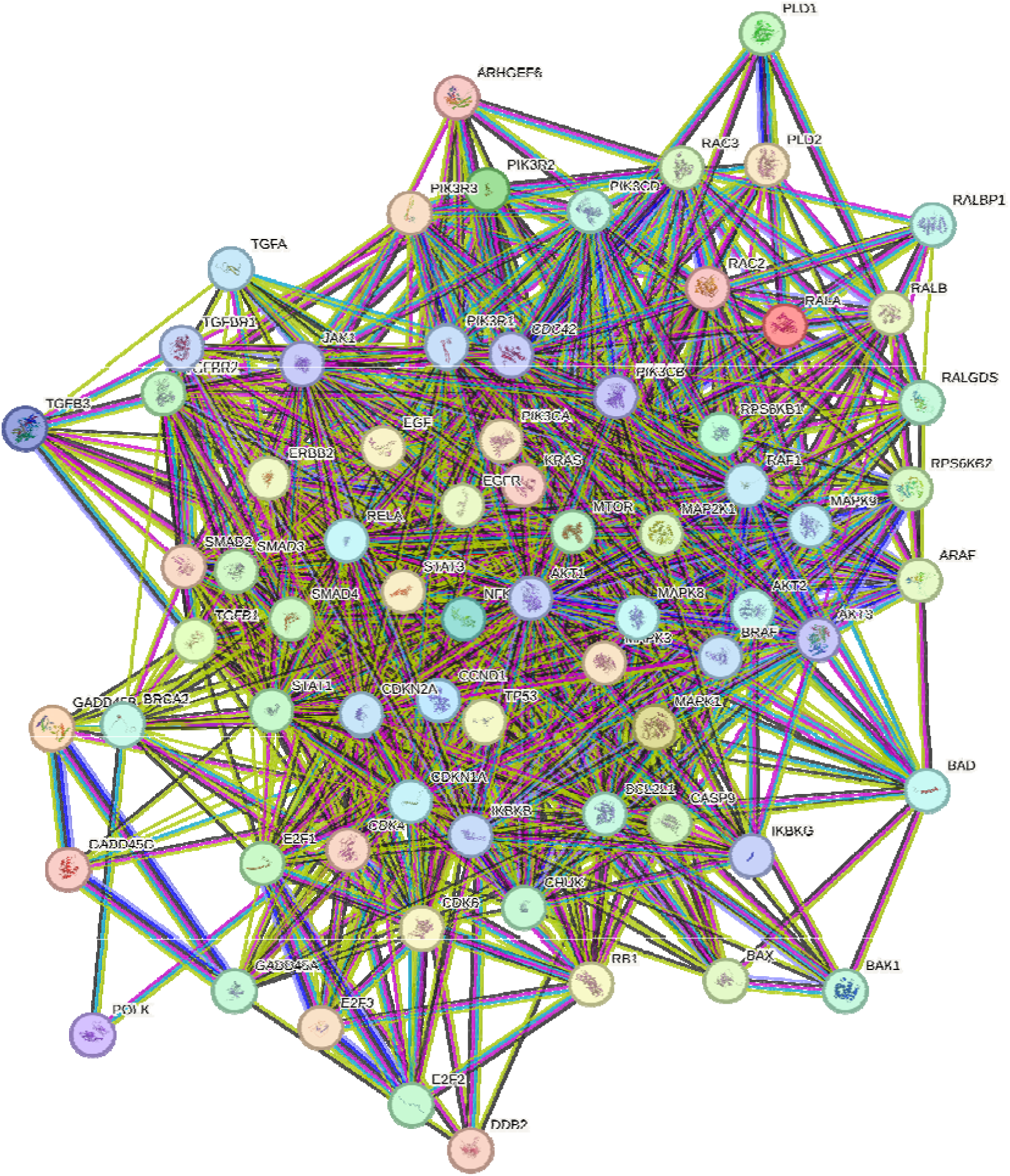
Network constructed by STRING utilizing the KEGG pathway HG0512. Content inside each node is known or predicted 3d structure of protein. Turquoise edges mean Protein-protein interactions from curated databases, purple means experimentally determined. Green, red, and dark blue edges indicate predicted Protein-protein interactions. Light green edges represent text mining, black represents co-expression, and light purple represents protein homology.

In table 3, we observe the ranking overlap for the top 50 genes for all three frameworks. 49 out of the top 50 highest ranked targets in GETgene-AI have been experimentally validated, while 47 were validated in STRING. However, only 30 of the highest ranking targets were validated in GEO2R.

**Table 3:**
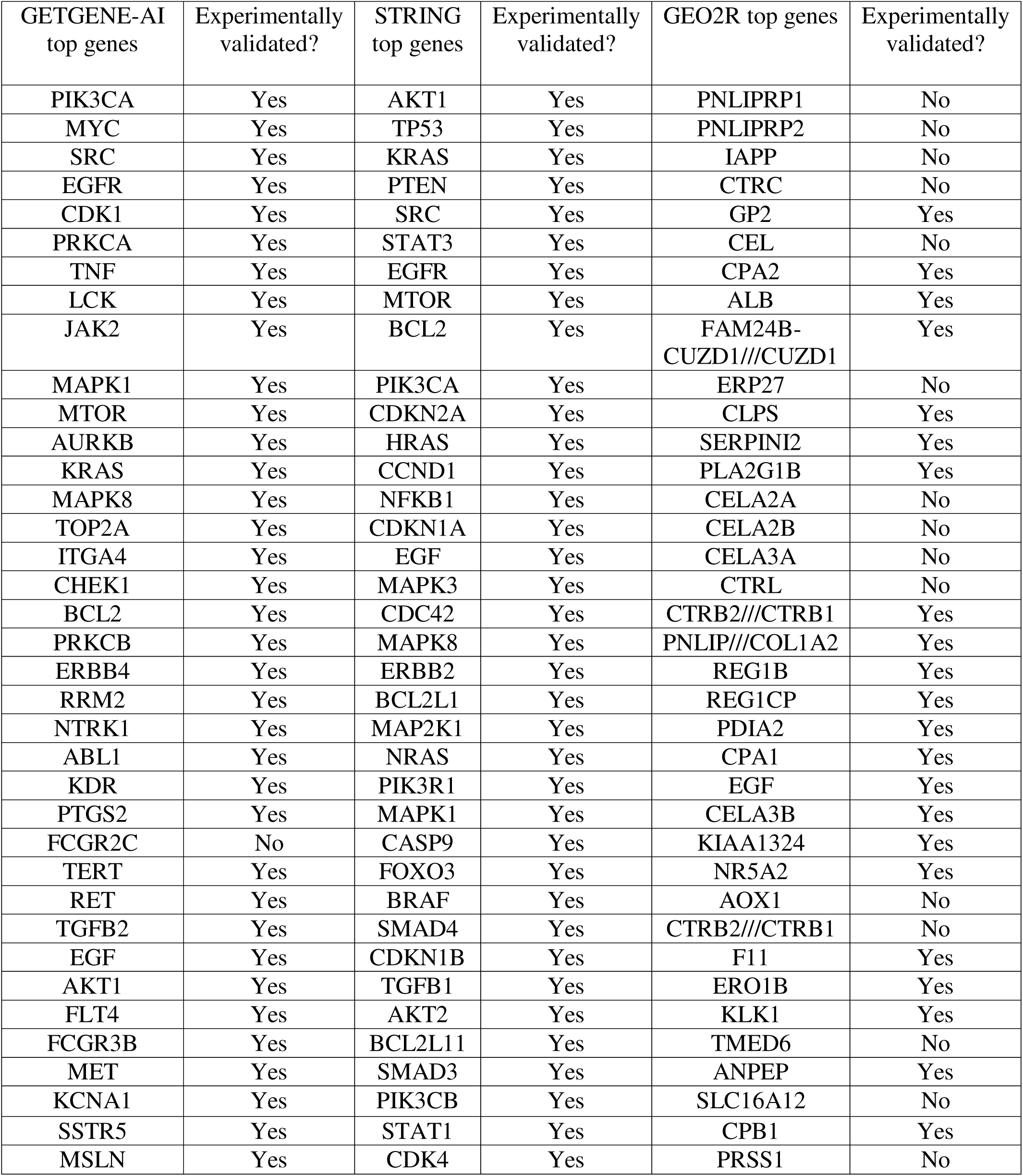
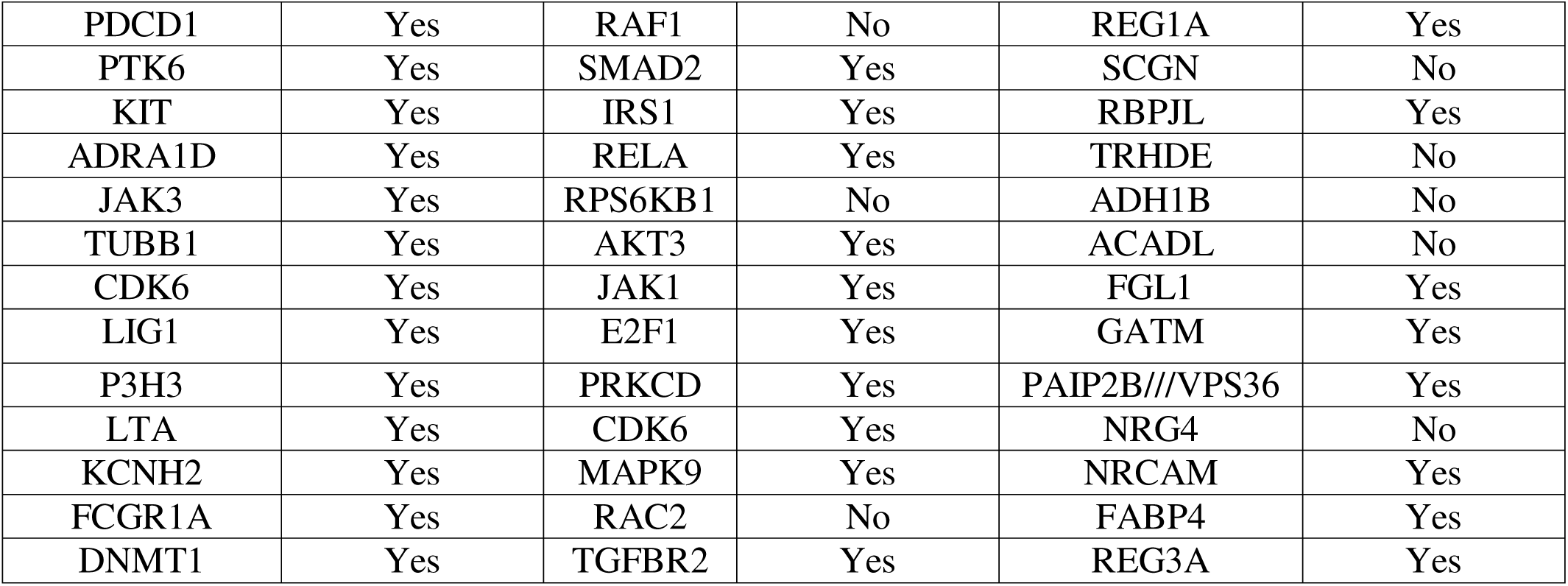
Top 50 genes from GETGENE-AI, STRING, and GEO2R and their status as experimentally validated drug targets.

Comparing GETgene-AI to GEO2R and STRING, our framework demonstrates a 38% improvement over GEO2R in the rate of experimental validation of the top 50 genes on each list. In figure 4, we observe the differences in the percentage of experimentally validated targets out of the top 50.

**Figure 4:**
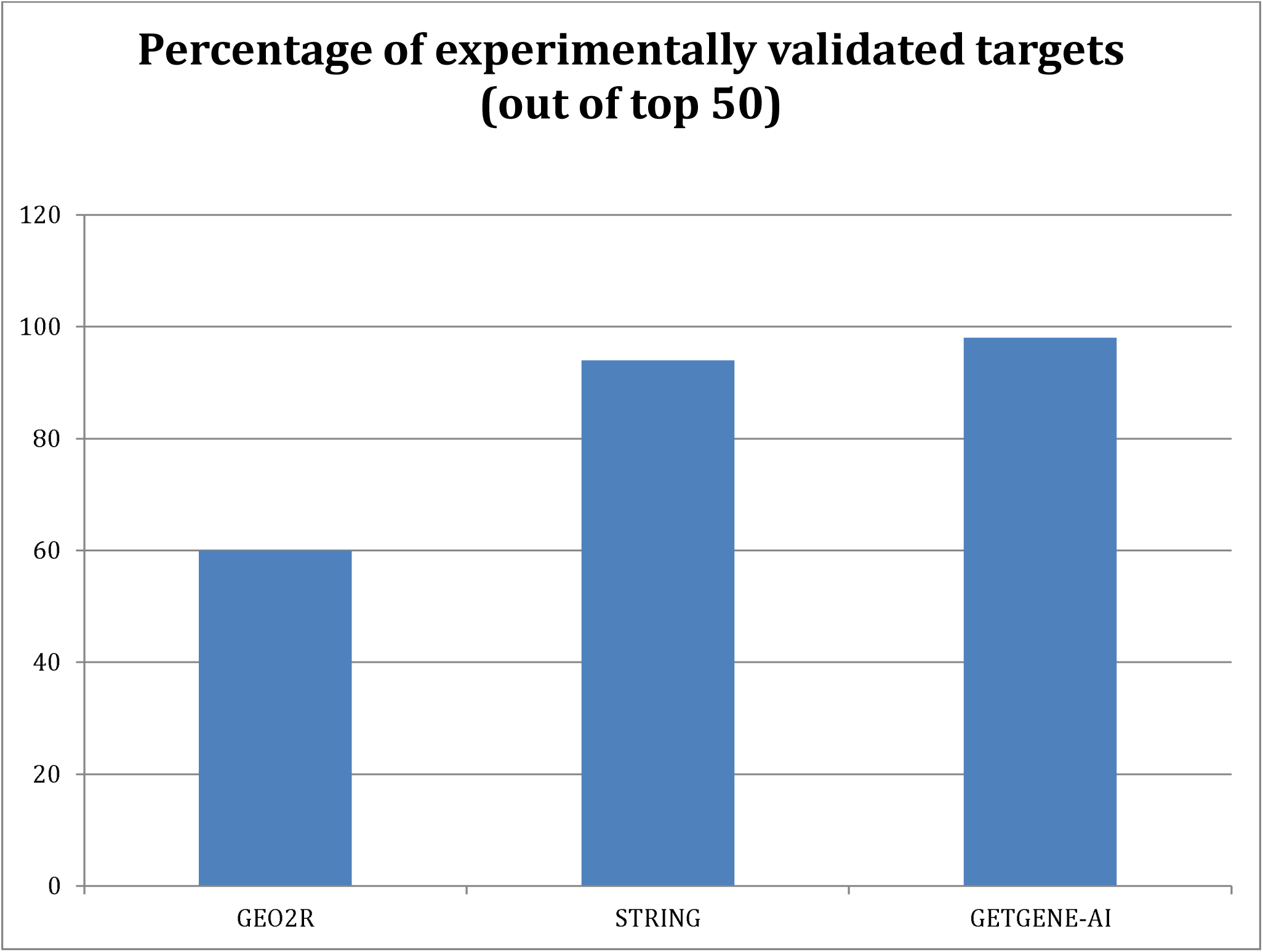
Bar graph displaying the percent of experimentally validated targets out of the top 50 genes with each framework.

### Enhancement provided by AI

GPT-4o was utilized to conduct a comprehensive literature assessment for our gene list. Although its output was not incorporated into the final weighted score, the GPT-4o scores demonstrated strong correlations with both the weighted score and all three GET list scores. Notably, GPT-4o prioritized genes such as *MYC* and *SRC*, reflecting their well-documented prominence in the scientific literature. This complemented GETgene-AI’s approach, which relies on network mutational analysis for gene prioritization. To minimize the inclusion of false positives in the GPT-4o scoring process, we instructed GPT-4o to directly cite articles from its internal database. While GPT-4o did not exhibit a higher rate of experimental validation compared to manual methods, it significantly reduced the time required for literature review by 80%. Human oversight was essential to ensure accuracy and reliability of GPT-4o in verifying the literature and reasoning of GPT-4o before incorporation into the pipeline.

The RP-LIT score and GPT-4o score showed a high degree of correlation, with extremely similar rankings for each gene. Based on Spearman correlation analysis, the GPT-4o score (out of 400) exhibited a correlation coefficient of +0.457 with the weighted score, indicating a statistically significant relationship. Table 4 provides a detailed comparison of the ranking differences between the GPT-4o score and the GET ranking score, highlighting the alignment and discrepancies between the two approaches.

**Table 4:** Top 20 highest ranked genes based off of GPT4 score compared to their ranks in GET and their status as experimentally validated drug targets.

| Gene | GPT4-Score<br>ranking | GET<br>ranking | Experimental<br>Validation? | Citation |
| --- | --- | --- | --- | --- |
| MYC | 1 | 2 | Yes | (Zhang et al., 2024) |
| SRC | 2 | 3 | Yes | (Su et al., 2023) |
| EGFR | 3 | 4 | Yes | (Wu et al., 2023) |
| TERT | 4 | 27 | Yes | (Campa et al., 2015) |
| RRM2 | 5 | 21 | Yes | (Li et al., 2022) |
| PIK3CA | 6 | 1 | Yes | (Payne et al., 2015) |
| TOP2A | 7 | 16 | Yes | (Pei et al., 2018) |
| NTRK1 | 8 | 22 | Yes | (Cheng et al., 2013) |
| PTGS2 | 9 | 25 | Yes | (Hingorani et al., 2003) |
| EGF | 10 | 30 | Yes | (Sheng et al., 2020) |
| CDK1 | 11 | 5 | Yes | (Huang et al., 2021) |
| MAPK1 | 12 | 10 | Yes | (Si et al., 2023) |
| KRAS | 13 | 13 | Yes | (Timar & Kashofer, 2020) |
| MTOR | 14 | 11 | Yes | (Stanciu et al., 2022) |
| MSLN | 15 | 37 | Yes | (Hu et al., 2024) |
| RET | 16 | 28 | Yes | (Bhamidipati et al., 2023) |
| AKT1 | 17 | 31 | Yes | (Arasanz et al., 2019) |
| JAK2 | 18 | 9 | Yes | (Huang et al., 2022) |
| MET | 19 | 34 | Yes | (Pothula et al., 2020) |
| PDCD1 | 20 | 38 | Yes | (Marabelle et al., 2020) |

### Broader Implications and Generalizability

While the current study focuses on pancreatic cancer, the GETgene-AI framework can be readily adapted to other cancers or diseases with access to similar genomic and clinical data resources. Future studies will explore its application to breast and lung cancers by employing the same systematic process described in this work. The GETgene-AI framework integrates literature review, large-scale sequencing data, and network centrality scores, providing a comprehensive approach to drug target prioritization. Additionally, its reliance on computational methods for prioritization and the elimination of statistically insignificant data ensures that the framework is both scalable and efficient, making it suitable for broader applications in biomedical research.

## DISCUSSION

Through the application of GETgene-AI to pancreatic cancer, we have identified several under researched promising drug targets: PIK3CA, PRKCA, LCK, MAPK8, ITGA4, PRKCB, and KCNA1. We have validated our hypothesis that the utilization of network-based analysis, artificial intelligence, and biologically significant data will enable systemic prioritization of actionable therapeutic targets. GETgene-AI’s approach to drug target prioritization integrates literature review, large-scale sequencing data, network-based centrality scoring, and assessment of potential adverse effects through organ expression scores. This multifaceted implementation offers a scalable and comprehensive framework for drug target prioritization, which can be readily adapted to other cancers with similar data availability. Additionally, the integration of an assessment of potential adverse effects through organ expression scores strengthens its applicability to finding targets with low adverse events in clinical trials. Furthermore, GETgene-AI’s ability to systematically deprioritize genes with low mutational relevance underscores its superiority in efficiently narrowing down actionable and biologically relevant targets. Slight variations of cutoffs utilized for the compilation and prioritization of the GET lists did not result in significant variations of the final rankings or scores of the final GETgene-AI gene list.

### False Positives and Limitations

False positives are an inherent risk in large-scale computational analyses. The GETgene- AI framework addresses this challenge through iterative refinement and the systematic exclusion of genes lacking functional or experimental support. Future validation efforts will focus on further refining these rankings through targeted experimental studies. Additionally, the literature assessment provided by generative AI is expected to improve as AI technology advances and our model is trained on more experimental data, thereby minimizing inaccuracies or "hallucinations" in the generated outputs.

To mitigate false positives, genes without functional relevance to cancer were systematically excluded. For instance, genes that ranked highly due to algorithmic artifacts but lacked experimental validation or literature support were deprioritized. Examples include *ITGA4* and *PRKCB*, both of which have fewer than 10 PubMed articles discussing their role in pancreatic cancer. These genes were ranked lower than many well-established targets due to their low scores in the GET, GT, and Expression lists, which prioritize targets with robust experimental or literature support during the RP score calculation process.

This study has several limitations. First, the top-ranked targets identified by GETgene-AI require further experimental validation, which is a critical next step to confirm their biological and therapeutic relevance. Second, the reliance on publicly available datasets may introduce biases due to incomplete or inconsistent annotations. These limitations highlight the need for further experimental validation and the incorporation of more comprehensive datasets to enhance the accuracy and reliability of the framework. Third, disease specific factors cannot be directly considered with GETgene-AI, such as protein aggregation in Alzheimer’s Disease (Pluta & Ułamek-Kozioł, 2020).

### Contributions and Limitations Provided by GPT4o

GPT-4o significantly enhanced the efficiency of literature-based ranking by helping automate the review and prioritization of scientific abstracts. This approach significantly increased the efficiency of literature review. However, inherent challenges, such as the risk of hallucination, necessitated manual verification to ensure the accuracy of the results. As such, artificial intelligence is utilized as a complementary tool for searching and assessing literature, rather than a replacement to human expertise (Blanco-González et al., 2023). While GPT-4o provides substantial value, its integration into research workflows should be approached cautiously, with safeguards implemented to mitigate potential errors. Additionally, training GPT- 4o on more experimental data in the future will further improve its accuracy and reliability in prioritization tasks.

### Future Directions

Future work will prioritize experimental validation of top-ranked targets, such as *PIK3CA* and *PRKCA*, using CRISPR-mediated knockouts in pancreatic cancer cell lines. Subsequent *in vitro* drug response assays will evaluate the therapeutic potential of these targets. Additionally, we aim to refine the framework by incorporating multi-omics datasets (e.g., proteomics, metabolomics) and enhancing its ability to predict adverse effects through improved organ expression profiling.ce of these targets.

While the current study focuses on cancer applications, future research will expand the scope of the GETgene-AI framework. We plan to validate its utility in additional cancer types, such as breast and lung cancer, and explore its applicability to non-cancerous disease contexts, including neurodegenerative disorders like Alzheimer’s and Parkinson’s.

## CONCLUSIONS

The GETGENE-AI framework represents a significant advancement in computational drug discovery, integrating network-based prioritization with machine learning to prioritize actionable therapeutic targets efficiently. Genes highlighted through our case study in pancreatic cancer such as *PRKCA*, *LCK*, *ITGA4*, and *PRKCB* are novel targets that require further exploration. While this study focuses on pancreatic cancer, the GETGENE-AI framework is adaptable to other cancers and diseases, offering a modular and versatile approach for target discovery. GPT4o enhanced the efficiency and accuracy of literature-based ranking, reducing manual workload and aligning well with network-based rankings. However, its reliance on manual verification underscores the need for cautious integration into automated pipelines. By refining target discovery methods, the GETGENE-AI framework paves the way for personalized therapeutic strategies and accelerates the translational research in oncology. Future work will focus on expanding the framework to other cancers, improving ranking metrics, and integrating multi-omnics datasets to enhance its predictive power. Future iterations of GETgene-AI aim to integrate multi-omics datasets, such as single-cell RNA-seq and metabolomics, to capture greater biological complexity.

## Data availability

All files containing genes and drugs utilized in the GETGENE-AI process have been uploaded to our Github repository https://github.com/alphamind-club/GETGENE-AI. “GEO2R assessment”, “GETGENE-AIvsGET”, “GETvsGPT4vsSTRINGvsGEO2R”, and “STRING Assessment” contain experimental validation assessments performed to compare the top 50 genes from each approach. Other files are the initial lists of genes utilized to form each of the GET lists before cutoffs were applied. ChatGPT’s GPT4o model was used for the GPT4o literature assessment score, which requires a monthly subscription. The custom GPT4o model made for this study is: Research GPT. ProteinAtlas was used to find the expression levels.

## Acknowledgment

Both authors thank the administrative support of AlphaMind Club for making this mentored research possible. JYC thanks the generous support of the startup fund to the Systems Pharmacology AI Research Center at UAB and NIH common fund grant award U54-OD036472, which partially supported this research. The authors acknowledge the use of ChatGPT in improving the structure and readability of the manuscript.

